# Automated conservation assessment of the orchid family using deep learning

**DOI:** 10.1101/2020.06.11.145557

**Authors:** Alexander Zizka, Daniele Silvestro, Pati Vitt, Tiffany M. Knight

## Abstract

IUCN Red List assessments are essential for prioritizing conservation needs but are resource-intensive and therefore only available for a fraction of global species richness. Tropical plant species are particularly under-represented on the IUCN Red List. Automated conservation assessments based on digitally available geographic occurrence records can be a rapid alternative, but it is unclear how reliable these assessments are. Here, we present automated conservation assessments for 13,910 species of the diverse and globally distributed Orchid family (Orchidaceae), based on a novel method using a deep neural network (IUC-NN), most of which (13,049) were previously unassessed by the IUCN Red List. We identified 4,342 (31.2 % of the evaluated orchid species) as *Possibly Threatened* with extinction (equivalent to the IUCN categories CR, EN, or VU) and point to Madagascar, East Africa, south-east Asia, and several oceanic islands as priority areas for orchid conservation. Furthermore, the Orchid family provides a model, to test the sensitivity of automated assessment methods to issues with data availability, data quality and geographic sampling bias. IUC-NN identified threatened species with an accuracy of 84.3%, with significantly lower geographic evaluation bias compared to the IUCN Red List, and was robust against low data availability and geographic errors in the input data. Overall, our results demonstrate that automated assessments have an important role to play in achieving goals of identifying the species that are at greatest risk of extinction.

## Introduction

Prioritizing the use of conservation resources is essential to counter the current global biodiversity crisis (Possingham et al. 2002; Rodrigues et al. 2006). The International Union for the Conservation of Nature (IUCN) global Red List (RL) is the most widely used scheme to evaluate species’ risk of extinction and is based on rigorous criteria and the best available scientific information (Rodrigues et al. 2006; Newton 2008; Pfab et al. 2011). The IUCN RL scores species’ population size, range size, population trends (decline or increase), and threats for a standardized assessment into extinction risk categories (Collen et al. 2016; IUCN Standards and Petitions Subcommittee 2017). Due to the rigor of the IUCN RL, they require intensive data, expertise and time (Roberts et al. 2016), and as a result, only a fraction of species are red-listed for many taxa (e.g., plants) and geographic areas (i.e., the tropics). For example, less than 20% of flowering plant species are IUCN red-listed (www.iucn.org; Troudet et al. 2017; Bachman et al. 2018).

The Orchidaceae is a diverse, globally important plant family with a high need for conservation assessment and prioritization (Swarts & Dixon 2017; Fay 2018). There are approximately 29,000 orchid species (Givnish et al. 2016) on all continents except Antarctica with a variety of life forms, from terrestrial to epiphytic (Cribb et al. 2003; Givnish et al. 2016). Many orchid species are local endemics and their distributions are constrained by edaphic environments and by their relationships with mycorrhizal fungi and specialist pollinators (McCormick & Jacquemyn 2014; Gaskett & Gallagher 2018). Orchids are economically important in horticulture and in the floral, pharmaceutical, and food industries (Subedi et al. 2013; Hinsley et al. 2018) and many species face immediate threats by land conversion and illegal harvesting (Hinsley et al. 2017; Fay 2018). Their global trade is so problematic that the Convention on Intentional Trade in Endangered Species of Wild Fauna and Flora (CITES) lists all orchid species with accepted names (Hinsley & Roberts 2018). Thus, there is an urgent need for identifying the most endangered species to prioritize *in situ* protection and *ex situ* conservation programs.

The need to prioritize species given limited data has triggered the development of methods for fast, automated conservation assessments (AA), based on digitally available species distribution data (for example from www.gbif.org). Two distinct classes of methods are available: 1) index-based methods that calculate multiple indices characterizing a species range size and use thresholds provided under IUCN Criterion B to classify species into IUCN RL categories. Multiple index-based methods exist (Bachman et al. 2011; Schmidt et al. 2017; Cardoso 2017; Dauby et al. 2017) and they can be used either to support the IUCN RL assessment process, or with additional assumption on habitat destruction and threat, to stand-alone as preliminary assessments (Schmidt et al. 2017; Cosiaux et al. 2018; Zizka et al. 2020c). 2) prediction-based methods that use existing IUCN RL assessments together with species traits to predict the conservation status of unevaluated or *Data Deficient* species (Bland et al. 2015; Pelletier et al. 2018; González-del-Pliego et al. 2019; Lughadha et al. 2019), including the use of machine learning algorithms. Prediction-based methods may use the same indices on species’ ranges as index-based methods, but also incorporate additional traits such as climatic niche, biomes, human footprint index, geographic region, or traits related to species morphology or physiology (Bland et al. 2015; Di Marco & Santini 2015).

While existing AA methods can separate threatened (IUCN RL categories *Critically Endangered, Endangered*, and *Vulnerable*) from non-threatened (*Near Threatened* and *Least Concern*) species with an accuracy between 80 and 95% in animals (Bland et al. 2015), a recent study suggests lower performance for Orchids, with an accuracy of 51-84% in a taxonomically and geographically limited sample of 116 species from New-Guinea, (Lughadha et al. 2019). A known issue with index-based AA, their dependency on data availability cause them to overestimate the extinction risk of species with few occurrence records available (Rivers et al. 2011). However, the dependency on data availability, remains untested for prediction-based methods. Furthermore, it is currently unknown how robust AA are to erroneous geographic input data and geographic sampling bias (Walker et al. 2020).

Here, we present AA for 13,910 Orchid species (47.3% of all known species) based on digitally available occurrence data and test the sensitivity of the results to method choice, data availability, errors in the geographic input data and sampling bias. Specifically, we address three questions:

1. How threatened are Orchids and where are the global centers of extinction risk for Orchid species? Based on the IUCN RL, we expect ∼30% of orchids to be identified as *Possibly Threatened* (www.iucn.org; Lughadha et al. 2019).
2. How accurate are different AA methods? Based on the existing smaller-scale studies we expect an accuracy between 60-90% (Lughadha et al. 2019) and a susceptibility to over-estimate the threat status of narrow-ranged species.
3. How robust are AA to limitations with data availability, variable data quality and geographic sampling biases? Digitally available species occurrence records are often biased towards certain geographic regions (Meyer et al. 2016; Zizka et al. 2020a) and contain erroneous or very imprecise coordinates (Zizka et al. 2019, 2020b). Therefore, we expect a bias of AA towards well-sampled geographic areas and life forms, and an increase in accuracy with geographic cleaning.

## Methods

### Orchid extinction risk

To address **question 1** we used the taxonomy of the World Checklist of Selected Plant Families (WCSP 2019), and obtained native records of Orchid species on August 26, 2019, at the Global Biodiversity Information Facility (www.gbif.org, 2019). We then obtained full IUCN assessments from www.iucn.org in August 2019 and standardized them into the five most relevant threat categories: *Critically Endangered* (*CR*), *Endangered* (*EN*), *Vulnerable* (*VU*), *Near Threatened* (*NT*), and *Least Concern* (*LC*), excluding species assessed under Red List versions older than 3.1, and species assessed as *Data Deficient* (*DD*).

We implemented a deep neural network algorithm (Goodfellow et al. 2016)—IUC-NN—in the Python (v. 3.7) program ruNNer (https://github.com/dsilvestro/ruNNer) using the TensorFlow (https://www.tensorflow.org) library to predict the conservation status of unassessed Orchid species. We based the predictions on four groups of features, derivable from digitally available occurrence records: *geographic* (mean latitude and longitude, longitudinal and latitudinal range, extent of occurrence (EOO), area of occupancy (AOO), number of locations (*sensu* IUCN) and occurrence records), *climatic* (the mean of 19 bioclim variables, Karger et al. 2017), *biome* (presence in 14 biomes; Olson et al. 2001) and *anthropogenic* (the mean human footprint index, Wildlife Conservation Society - WCS & International Earth Science Information Network - CIESIN - Columbia University 2005).

We trained IUC-NN on all species with an IUCN RL assessment and occurrence records available. Prior to the training, we randomly split the dataset into a training set (90% of the entries) and a test set (10%). To carry out the training we further split the training dataset in 80% of the entries for training and 20% for validation. As the size of the dataset was comparatively small, we performed cross-validation by shifting the validation set five times to quantify the average validation cross-entropy loss and accuracy. We then used the neural network yielding the lowest cross-entropy loss across a range of models with different numbers of hidden layers and subsets of features to predict the conservation status of all Orchid species at two levels: binary (*Possibly Threatened* vs. *Not Threatened*) and detailed (*CR, EN, VU, NT*, and *LC*). See Supplementary Material S1 for details on the network architecture and training. We combined the IUC-NN assessments with distribution data from WCSP, to show the number and proportion of *Possibly Threatened* species and of evaluated species per geographic region (TDWG-level 3).

Since different habitats warrant different conservation approaches, we also summarized the number of orchid species and the fraction of threatened species in major biomes of the world. We classified species into biomes using the ‘speciesgeocodeR’ v. 2.0-10 package (Töpel et al. 2017) if at least five percent of a species’ records occurred in this biome.

### The accuracy of AA

To address **question 2**, we performed AA with three index-based methods: ConR v.1.2.2 (Dauby et al. 2017), speciesgeocodeR v.2.0-10 (Schmidt et al. 2017; Töpel et al. 2017) and rCAT v.0.1.5 (Moat 2017) for all species with an IUCN RL status and occurrence records available. ConR, speciesgeocodeR and rCAT calculate EOO and AOO based on geographic occurrences and use thresholds of IUCN Criterion B to classify species into IUCN RL categories. Additionally, ConR calculates the number of locations at which a species occurs. We then compared the accuracy of all four AA methods on both the binary and detailed levels with the existing IUCN RL as standard. We provide full error matrices for both levels of assessment. For the binary assessment, we also calculated the fraction of false positives (non-threatened species classified as *Possibly Threatened* by AA) and false negatives (threatened species classified as *Not Threatened* by AA). See Supplementary Material S1 for details on the index-based AA. Because the IUCN RL is biased both geographically and taxonomically (www.iucn.org), we repeated the accuracy tests of ConR with four additional reference datasets: (1) a random sub sample of IUCN RL (the “Sampled Red List index”, SRLI, Royal Botanic Gardens 2010); (2) species IUCN RL assessed after 2008 (IUCN2008), (3) species with at least 15 occurrence records available for AA (IUCN15); and (4) global conservation assessments from the ThreatSearch literature database. We only tested these alternative references with ConR, since we expect the other index-based methods to have similar results, and the alternative datasets included too few species to train IUC-NN models.

### The reliability of AA

To address **question 3**, we tested how the accuracy of IUC-NN dependent on the number of occurrence records available per species, different levels of geographic cleaning of the input data and the evenness of geographic sampling. See Supplementary Material S1 for details on the individual tests.

*Data availability* We used binomial regressions to test the dependence of AA and IUCN RL agreement on the number of occurrence records available for a species, using the stats::glm function in R, with a logit link.

*Data quality* We compared the accuracy of ConR assessments based on different datasets of species occurrences, representing three levels of data curation (1) “raw”, the data downloaded from GBIF scrubbed taxonomically, (2) “intermediate”, the raw data subjected to automated removal of records with common geographic errors (Zizka et al. 2020b), and (3) “filtered”, the intermediate data with additional removal of records outside the known occurrence range of species from WCSP (WCSP 2019). We only ran this test with ConR, since we expect the two other index-based methods to respond similarly to the issue and because we expect IUC-NN to be more robust against erroneous coordinates, since the features used for IUC-NN prediction are mean values across all records of a species.

*Sampling bias* Since IUCN RL and digitally available occurrence records are geographically biased (Meyer et al. 2016; Daru et al. 2018; Zizka et al. 2020a), we tested whether the observed differences in the statistical distribution of species over growth form types and geographic regions among (1) all orchid species, (2) IUCN RL assessed species and (3) species in our AA were greater than expected by random chance based on a null model in which we randomly permuted bias measurements among model types.

We did all analyses in R (R Core Team 2019) and Python, the analysed data and scripts, including the trained IUC-NN models will be available upon publication of this manuscript at a zenodo repository (10.5281/zenodo.3862199).

## Results

### Orchid extinction risk

We obtained 3,321,927 records with complete names for 19,034 taxa from GBIF. Of those, we retained 3,050,875/17,971 (records/species) for the “raw” dataset, 1,188,658/16,935 for the “intermediate” dataset, and 999,476/14,148 for the “filtered” dataset (see Supplementary Table S1 in Supplementary Material S2 for the effect of individual filters). We obtained IUCN RL assessments for 1,404 species, and retained 861 species (49.7% of them *Threatened*) fitting our criteria to train the IUC-NN neural network.

We generated IUC-NN assessments 13,910 species (47.3 % of all species in the family; Supplementary Material S3). On the binary level, 9,772 (68.9%) classified as *Not Threatened* and 4,415 (31.1%) classified as *Possibly Threatened*. On the detailed level 10,733 species (75.7%) classified as *LC*, 51 species (0.4%) as *NT*, 70 species (0.5%) as *VU*, 2,610 species (18.4%) as *EN*, and 723 species (5.1%) as *CR*. As the accuracy of the binary assessments was higher, we focus the visualization of our results and discussion on these, but provide results for the detailed assessment in the supporting information.

We identified three continental-level centers for Orchid conservation: northern South America, East Africa and south-east Asia with high numbers and proportions of *Possibly Threatened* orchid species (Fig. 1a). The regions with the most *Possibly Threatened* species were: Madagascar (646 species), Colombia (427), Borneo (369), and China South-Central (364) (Fig. 1a). Of those areas with more than ten Orchid species, the twelve most relevant regions included 6 islands: Réunion (73% of species *Possibly Threatened*), Mauritius (62%), Comoros (56%), Taiwan (51%), Seychelles (50%), and the Christmas Islands (50%) as well as 6 continental regions: Madagascar (86%), China South-Central (58%), Tibet (54%), Nepal (50%), Vietnam (49%), and East Himalaya (48%) (Fig. 1b). See Supplementary Figure S1, for a map of the detailed assessment.

**Figure 1:**
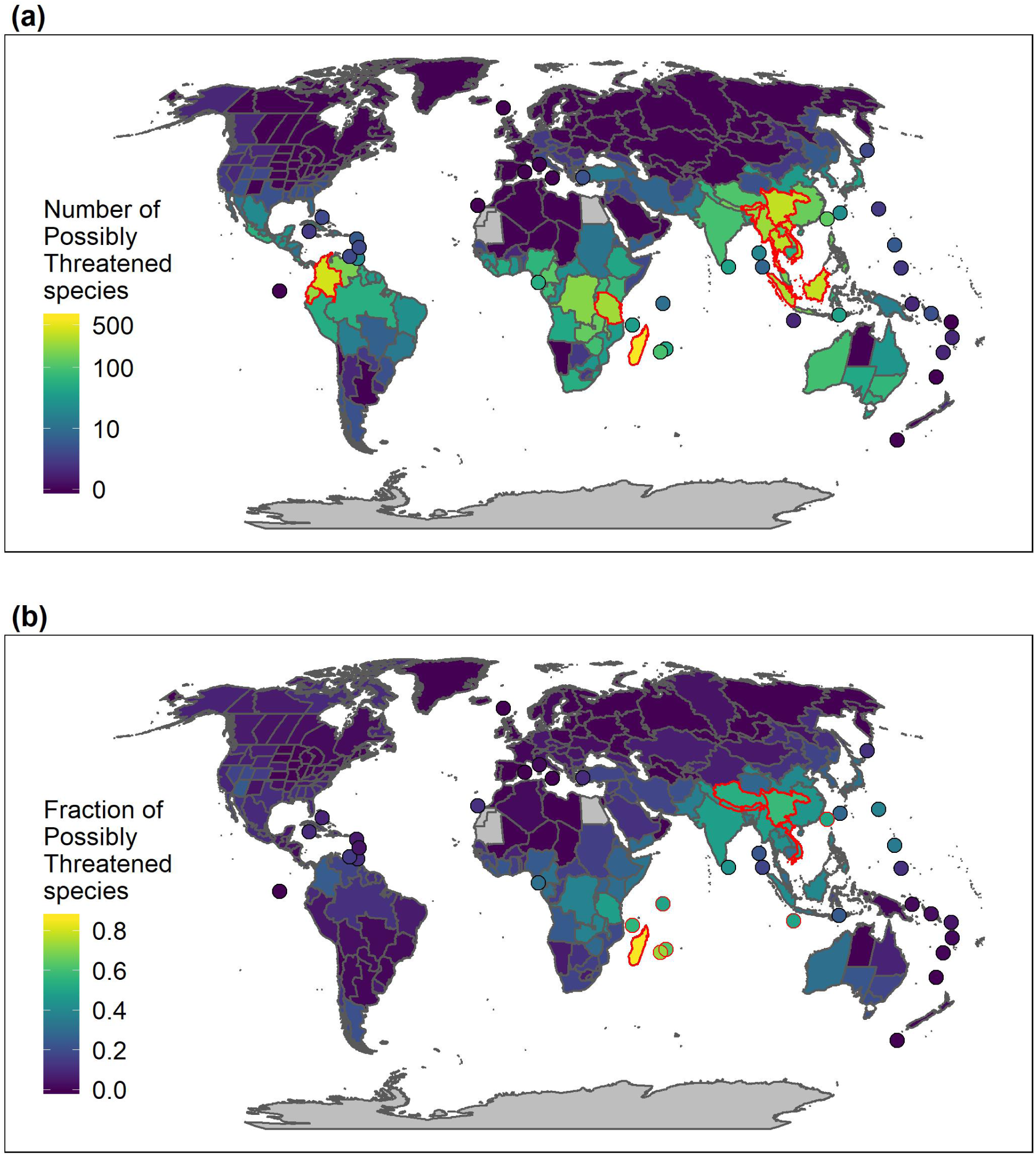
Global conservation status of orchids. **(a)** The number of possibly threatened orchid species per TDWG region. **(b)** The proportion of possibly threatened Orchid species on the number of evaluated species. Small islands and archipelagos with more than 10 known Orchid species are shown as points for better visualization. The red outlines indicate the 12 areas with the highest number (in a) and highest fraction (in b) of *Possibly Threatened* species. Region refer to TDWG level 1.

The number of *Possibly Threatened* Orchid species was highest in the Tropical moist broadleaved forest biome (3254 species) following the general diversity pattern of Orchids (Fig. 2, see Supplementary Figure S2, for the detailed assessment). In contrast, the proportion of *Possibly Threatened* Orchid species was highest in the Temperate conifer forest biome (45.8% of 365 species).

**Figure 2:**
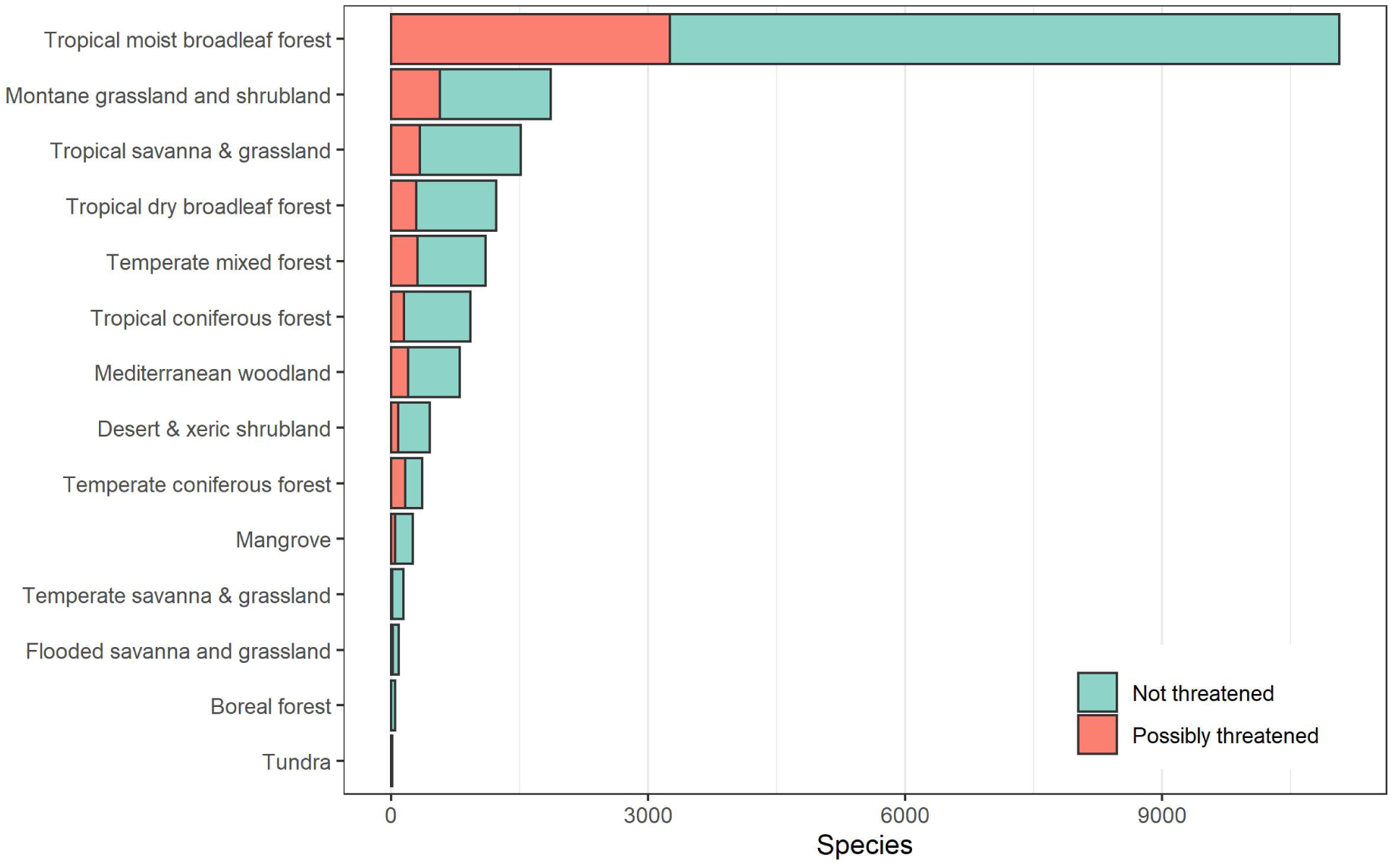
The threat status of Orchid species in the biomes of the world (Olson et al., 2001). Tropical moist broadleaf forest was the most specious habitat, and harbored the highest number of *Possibly Threatened* orchid species. The proportion of *Possibly Threatened* species was highest in temperate conifer forest.

IUC-NN increased the proportion of species evaluated across regions by a median of 53% (Fig. 3), with the highest increase in continental areas (TDWG level 3) in Gambia (from 0% of species evaluated to 100%), Southern Australia (from 1% to 94%), and South Argentina (from 0% to 92%).

**Figure 3:**
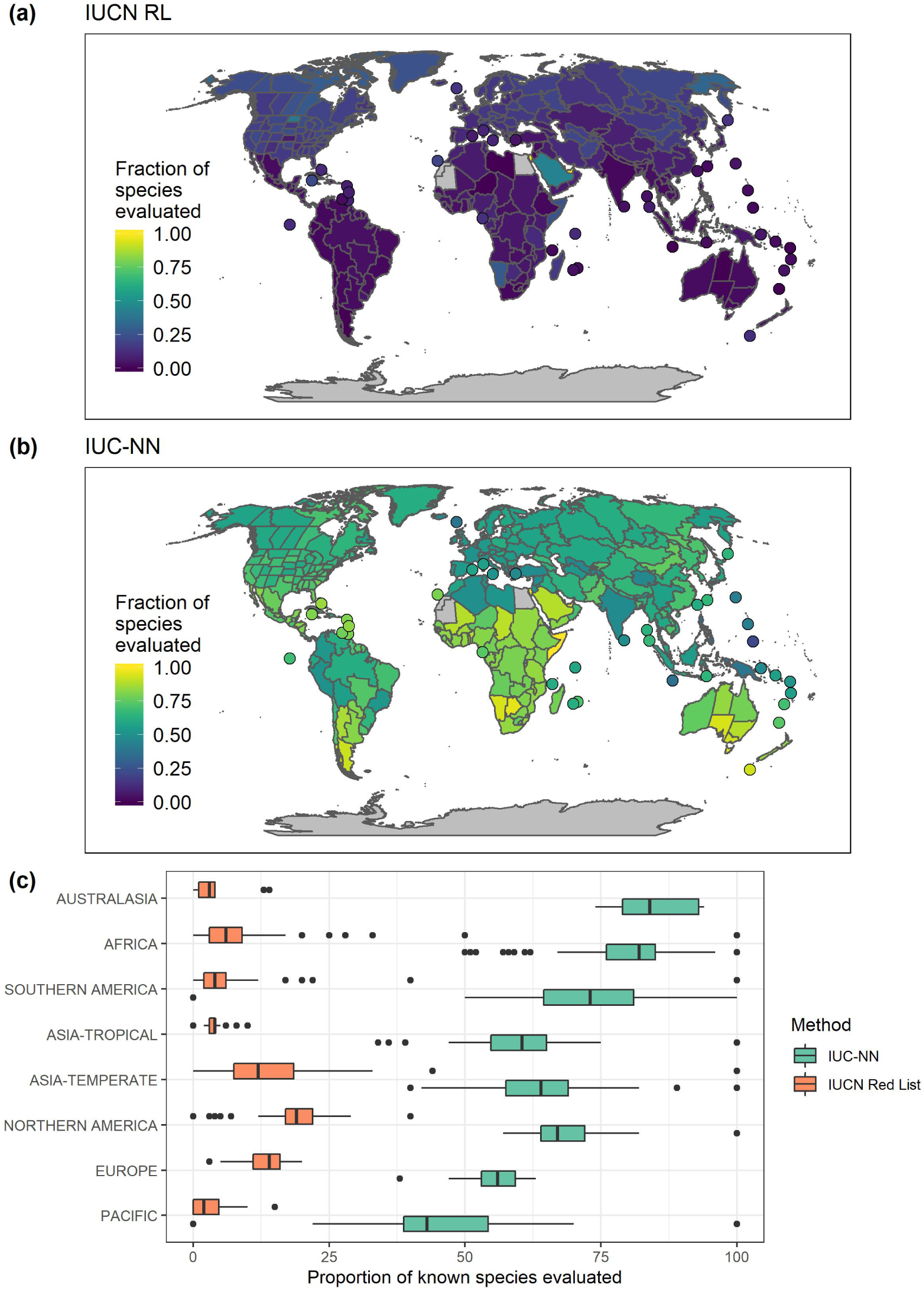
The fraction of Orchid species per geographic region evaluated for their conservation status. **(a)** IUCN Red List, **(b)** IUC-NN (prediction-based automated conservation assessment). **(c)** The proportion of species evaluated on the continental scale under both schemes. Each data point shows a geographic (TDWG) region and the y-axis indicates the number of percentage points by which IUC-NN increased the fraction of species evaluated.

### The accuracy of AA

The accuracy of all tested AA methods varied between 44.3% and 84.3%. As expected, the accuracy was higher separating *Possibly Threatened* from *Not Threatened* species (65.5% - 84.3%, Fig. 4b) as compared to the detailed assessments (44.3% - 64%, Fig. 4a, Supplementary Tables S2 and S3). At both levels the accuracy was similar for the index-based methods but higher for IUC-NN (Fig. 4).

**Figure 4:**
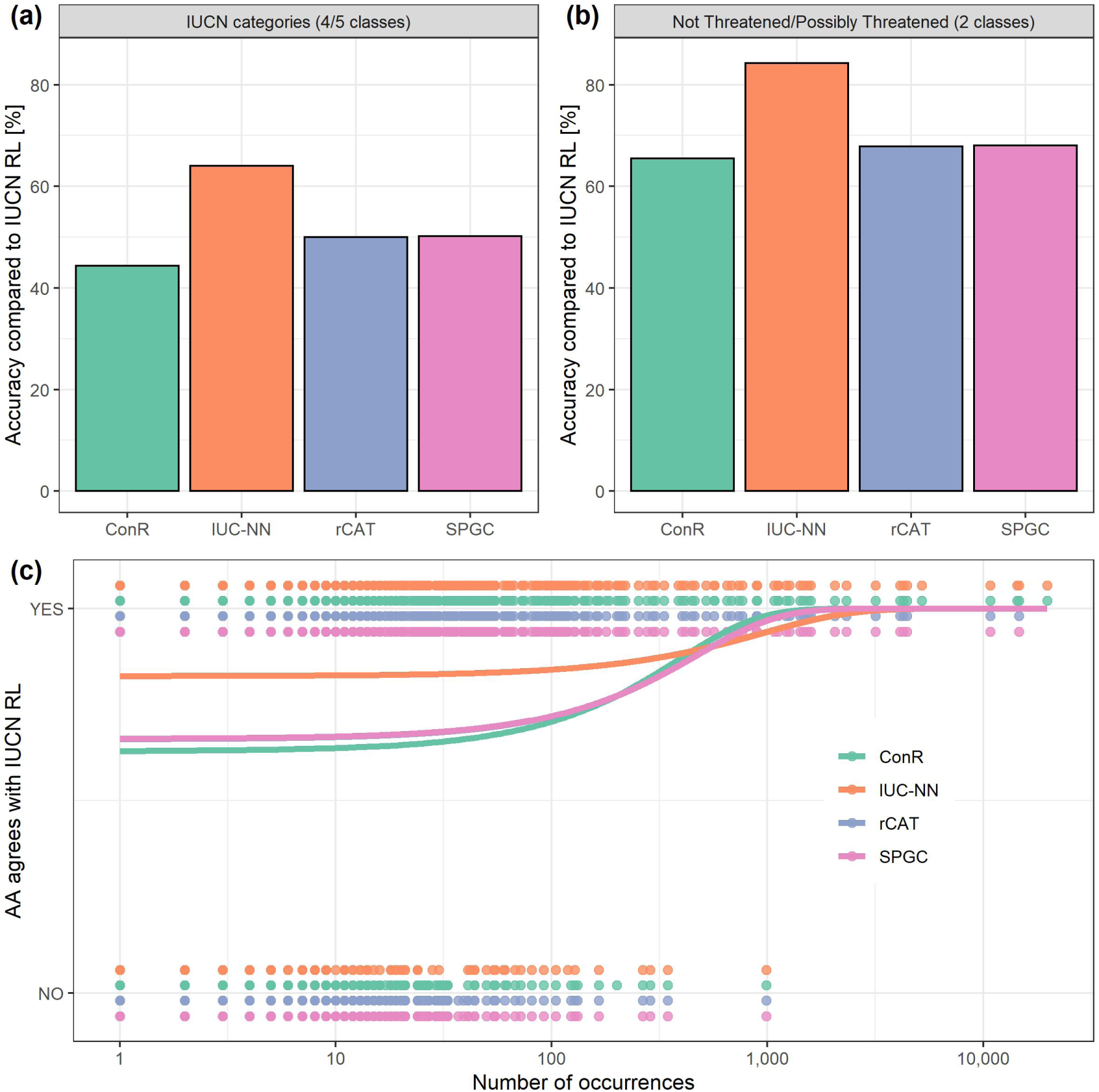
The accuracy of different methods for automated conservation assessment compared to the IUCN RL as gold standard. **(a)** The accuracy when assessing species into full IUCN categories (*CR, EN, VU, NT, LC*, the last two combined for the index-based methods) **(b)** The accuracy to identify species as *Possibly Threatened* or *Not Threatened* **(c)** The probability of classifying species accurately as *Possibly Threatened* or *Not Threatened* depending on the number of occurrence records available per species (n = 861). Not the logarithmic x-axis. In (c) the curves for rCAT and SPGC are identical.

The best IUC-NN model considered binary threat levels, used only geographic features and reached a test accuracy of 84.3% (88% positive predictive value and 79% negative predictive value; Table 1). Adding other features decreased the validation accuracy, but still remained within ∼5% of the best model (Supplementary Material S4). The best IUC-NN model on the detailed level included geographic features and the human footprint index and reached a test accuracy of 64% (Supplementary Material S4). The accuracy was generally higher for *CR* and *EN* as well as *LC* classes but low for the intermediate *VU* and *NT* classes (Supplementary Material S2). Furthermore, the accuracy was highest for species the IUCN considered threatened by “Natural systems modification” and “Energy production and mining”, and the lowest for species threatened by “Human intrusion & disturbance” and “Pollution” (Supplementary Figure S3). Concerning alternative reference datasets as gold standard instead of IUCN RL, the binary accuracy of index-based methods increased when using IUCN15 (69.9%, n = 359) or ThreatSearch (78.6%, n = 14), was similar for IUCN2008 (64.4%, n = 810) and lower for SRLI (57.1%, n = 261, Supplementary Table S4).

**Table 1:** Confusion matrices for different automated assessments in comparison to the IUCN RL assessment. For the index-based methods the results use all species with digitally available occurrence records (n = 866), for IUC-NN results from the test dataset are shown (n = 89).

| AA method | Spatial<br>cleaning | IUCN Assessment |  | Type |
| --- | --- | --- | --- | --- |
|  |  | Not<br>Threatened | Possibly<br>Threatened |  |
| **ConR** |  |  |  |  |
| Not Threatened | full | 75.1 | 24.9 | Index-based |
| Possibly Threatened | full | 38.8 | 61.2 | Index-based |
| Not Threatened | medium | 75.1 | 24.9 | Index-based |
| Possibly Threatened | medium | 36.4 | 63.6 | Index-based |
| Not Threatened | raw | 73.8 | 26.2 | Index-based |
| Possibly Threatened | raw | 35.3 | 64.7 | Index-based |
| **rCat** |  |  |  |  |
| Not Threatened | full | 68.3 | 31.7 | Index-based |
| Possibly Threatened | full | 32.5 | 67.5 | Index-based |
| **SPGC** |  |  |  |  |
| Not Threatened | full | 68.5 | 31.5 | Index-based |
| Possibly Threatened | full | 32.4 | 67.6 | Index-based |
| **IUC-NN best model** |  |  |  |  |
| Not Threatened | full | 78.9 | 21.1 | Prediction-based |
| Possibly Threatened | full | 11.8 | 88.2 | Prediction-based |
| **IUC-NN all features** |  |  |  |  |
| Not Threatened | full | 81.6 | 18.4 | Prediction-based |
| Possibly Threatened | full | 29.4 | 70.6 | Prediction-based |

The training and cross-validation of the IUC-NN model took less than 24 hours and the IUC-NN assessments for all species using the trained network took about 0.0005 seconds per species on a standard laptop. For the index-based methods, the median computation time was 0.02 seconds per species for rCAT, 0.13 seconds for SPGC, and 0.31 seconds for ConR on a standard laptop (n = 945), but increased with the number of records, in particular for ConR (Supplementary Figure S4) up to a maximum of 33.3 hours for *Gymnadenia conopsea* with 31,055 records.

### The reliability of AA

*Data availability* The accuracy of all index-based methods depended significantly on the number of records available (rCAT & SPGC: p=0.016, ConR: p=0.0056, n=861), but that was not the case for IUC-NN (p=0.14, n=861, Fig. 4c).

*Data quality* The geographic filtering of the input data only marginally affected by the accuracy of ConR, which varied between 65.5 - 68% among methods for the “filtered” dataset, 67.1 - 68.7% for the “intermediate” and 67.3 - 67.4% for the “raw” dataset.

*Sampling bias* The sampling intensity of the geographic input data was highest in Central and Northern Europe, Southern Australia, and Northern and Central America and was low in south-east Asia (Supplementary Figure S5). We found a difference in the evaluation frequency between the IUCN RL and AA based on life form and geography. Terrestrial species were over-represented on the IUCN RL (Fig. 5a), as were species from Africa, temperate Asia and Europe (Fig. 5b). Despite the strong bias in the availability of species occurrence data, we found significantly lower evaluation bias for AA than for the IUCN RL with respect to life form (epiphytic, lithophytic, terrestrial, mixed; p = 0.00465) and geographic region (p = 0.01855, see Supplementary Figure S6 for the interaction).

**Figure 5:**
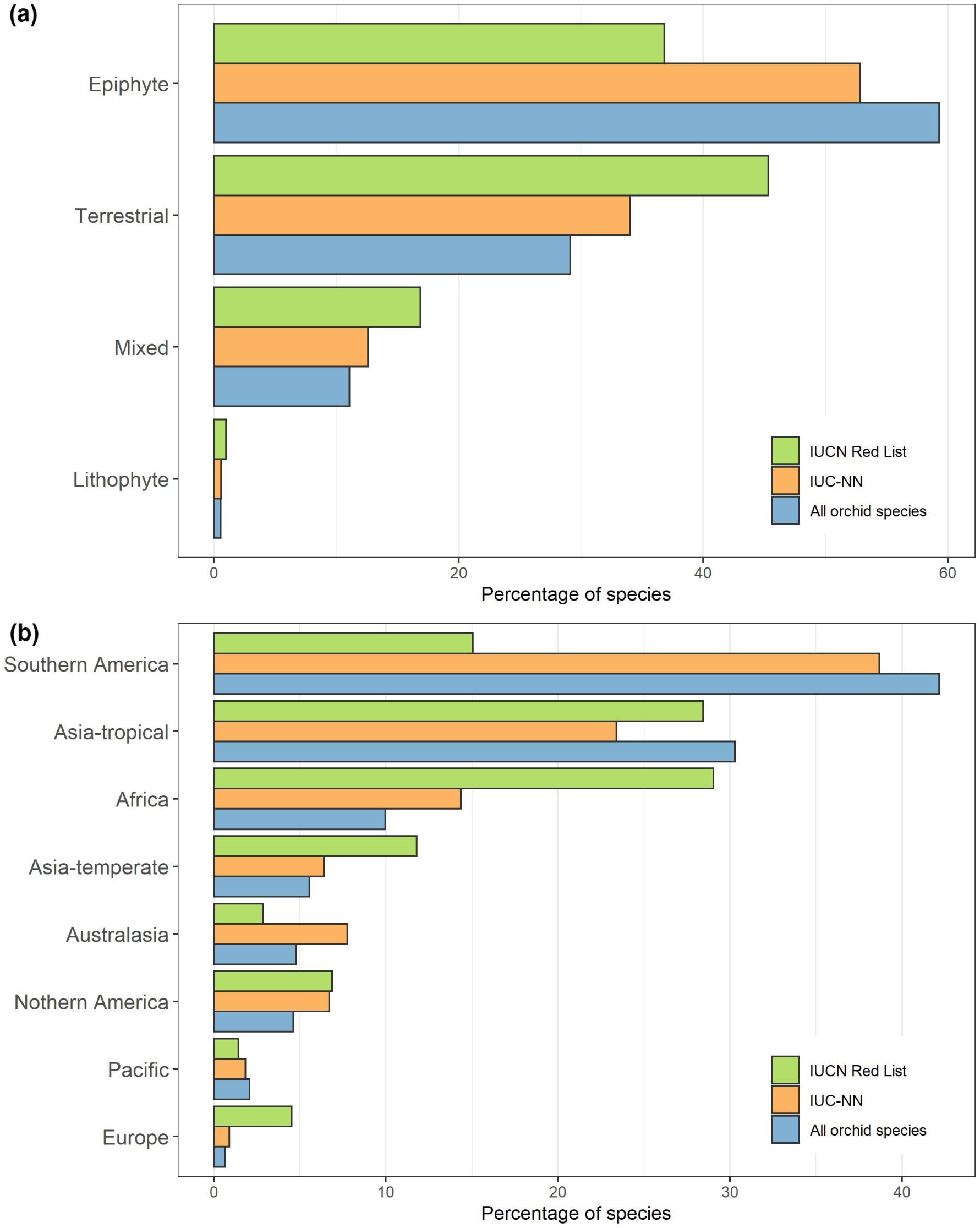
The bias in the conservation assessment of Orchids. **(a)** Life form. Most orchids with a known life form are epiphytes, but terrestrial species are over-represented in the IUCN Red List, potentially due to accessibility and geographic distribution. The automated assessment based on distribution records reduced this bias. **(b)** Geography. Most orchid species occur in South America and tropical Asia, but the IUCN Red List is biased towards Africa, temperate Asia and Europe. The automated conservation assessment reduced this bias. Regions refer to TDWG level 1.

## Discussion

### Orchid extinction risk

We found that 31.1% (4,415) of the Orchid species we were able to assess are *Possibly Threatened*, which is comparable to the recent estimate of 29.5% of global land plant species (Pelletier et al. 2018). As we could only evaluate species with available occurrence information, it is likely that an even higher proportion of those species we were unable to evaluate are *Possibly Threatened*. Indeed, unsampled species are likely to be rare and therefore infrequently encountered and identified. This possibility, combined with information on the generally high pressure humans have on orchids (Fay 2018), confirms that there is no doubt that the Orchidaceae family is in need of immediate conservation prioritization. Our findings also highlight that the risk is not equally distributed across their life forms, biomes and geographic regions.

We found that the distribution patterns of *Possibly Threatened* Orchids are related to total orchid species richness. There are high numbers of *Possibly Threatened* species in the species-rich tropics, especially the Neotropics, which are an important center of diversification for the Orchids with more than 11,000 Orchid species (Givnish et al. 2016), and in the Oriental and Papua-Melanesian biogeographic regions. These centers of orchid extinction risk coincide well with known areas of endemism, and hence small range sizes, for all plants (Kier et al. 2009). The North American Orchid flora is surprisingly depauperate given its large area (Gaskett & Gallagher 2018), and we found that few species in this region are *Possibly Threatened* (Fig. 1, Fig. 2). A similar pattern exists for northern Europe and Asia, although there are some *Possibly Threatened* species in the north of Asia. This general pattern may be associated with the climate of these north temperate regions, concomitant with a high proportion of terrestrial species.

It is well established that many species in the Orchid family need conservation attention (Dixon 2003; Wraith & Pickering 2018; Fay 2018; Liu et al. 2020), and there is an abundance of expertise and information available to successfully implement conservation strategies for both terrestrial (Swarts & Dixon 2009, 2017) and epiphytic species (Cribb et al. 2003; Swarts & Dixon 2017; Fay 2018). Most species are facing more than one threat and have limited distributions (Wraith & Pickering 2018). Until now, what has been lacking is a comprehensive assessment of the conservation status this large and highly diverse family, particularly for understudied species and geographic regions.

### The benefits and challenges of AA

The 84% accuracy of IUC-NN to correctly identify *Possibly Threatened* species suggests that it can be used with confidence for conservation science and practice. The major advantages of IUC-NN are that it is: 1) orders of magnitude faster than full IUCN RL assessments (Supplementary Figure S4), 2) objective and reproducible, 3) based solely on digitally available data, and 4) able to incorporate data beyond the formal IUCN RL Criteria (Bland et al. 2015). Furthermore, IUC-NN is less sensitive than index-based methods to limited available occurrence data and displayed a lower false positive rate (Table 1, Fig. 4; Rivers et al. 2011; Lughadha et al. 2019).

The overall accuracy of IUC-NN was similar to the performance of prediction-based methods in other taxa (e.g. amphibians, González-del-Pliego et al. 2019; mammals, Bland et al. 2015; non-orchid monocots, Darrah et al. 2017 and land plants @Lughadha2019). However, the mis-classification rate of ∼15% is undesirable when making specific conservation decisions. A close examination of the species which IUC-NN misclassified most severely (wrong by the binary model and wrong by the maximum number of categories by the detailed model; n = 11; Supplementary Material S5), provides insight on the main causes of misclassification. The four species for which IUC-NN overestimated extinction risk: *Ancistrorhynchus parviflorus, Disperis elaphoceras, Oligophyton drummondii, Tridactyle stevartiana* all are known from only a few locations in East Africa, and were assessed as *LC* on the IUCN RL because these locations are within protected areas, on the strong assumption that these guarantee effective and long-term protection. The seven species IUC-NN underestimated extinction risk: *Anguloa cliftonii, Brachionidium pteroglossum, Bulbophyllum sceliphron, Caladenia hastata, Paphiopedilum schoseri, Paphiopedilum supardii, Vanilla cribbiana*, the IUCN RL assessment included information regarding human changes to the local environment, severe collection pressure for the horticultural trade, or observed population declines. In particular, the terrestrial orchid *C. hastata* is already the focus of active recovery (www.iucn.org), and *B. sceliphron* might be even more threatened than it is currently listed and is in need of further assessment according to the detailed IUCN RL assessment.

Including data on species trade, land use or life history, once they are available, may help to increase the accuracy of IUC-NN and to overcome the dependency on threat category (Supplementary Figure S3). Further, combining the binary and detailed model, which contradicted in some cases (Supplementary File 4), might increase the performance of IUC-NN. We suggest that labelling species where the binary and detailed model disagree as *Data Deficient* might reduce the number of false predictions. Future developments of IUC-NN will quantify the estimation of threat as a continuous parameter to increase the accuracy for intermediate threat levels and will implement Bayesian neural networks to quantify uncertainties in the prediction (Silvestro & Andermann 2020).

Our results suggest that AA are robust to geographic errors in the input data and reduce evaluation bias compared to the IUCN RL (Fig. 5). Although AA are certainly not free from error, due to the biased availability of the underlying occurrence records and RL data (Meyer et al. 2016, Supplementary Figure S5; Daru et al. 2018), the reduced bias compared to IUCN RL is likely related to the fact that the factors biasing the collection of species geographic occurrence records, such as physical accessibility or socio-economic factors (Meyer et al. 2016; Daru et al. 2018; Zizka et al. 2020d), affect the collection of detailed data necessary for IUCN RL assessments (e.g., population trends) even more strongly.

Cleaning spatial data did surprisingly little to improve the accuracy of AA (Table 1), considering that geographic errors can erroneously increase the range size of species by thousands of kilometers. This result is likely due to low number records available for many Orchid species. In such a setting, the reduction of range size produced via the removal of erroneous points might have improved the estimation of the range shape, but not the range size, as the range size is already underestimated for most species.

The advantages of IUC-NN come at the price of applying complex statistical machinery and moving beyond strict adherence of IUCN criteria, which warrants some caution (Walker et al. 2020). We see the major application of IUC-NN in integrating diverse types of data into preliminary conservation assessments identifying species or areas in need of protection or more detailed assessment (Fig. 1). This approach is particularly valuable for the species-rich, but data-poor, tropical regions of Earth (Mounce et al. 2018), where even preliminary assessments will be useful for informing conservation. A particular strength of IUC-NN is that it can be trained for other taxonomic groups or regions contributing to speeding up the conservation assessment of all species on Earth.

## Conclusions

We classified 4,342 orchid species (31.2% of the family) as *Possibly Threatened*, or on a more detailed scale 718 (5.2%) as *Critically Endangered*, 2,567 (18.5%) as *Endangered*, and 68 (0.5%) as *Vulnerable* (question 1). Northern South America, Madagascar and south-east Asia as well as the Tropical moist broad leaf forest biome harbor the highest number of *Possibly Threatened* species. In relation to total species richness we identified Madagascar, oceanic islands as well as parts of East Asia as the centers of *Possibly Threatened* species.

We showed that the IUC-NN method based on digitally available data can identify *Potentially Threatened* Orchid species with an accuracy of over 84%, and individual IUCN categories with an accuracy of 64% (question 2). IUC-NN provides preliminary conservation assessments for 13,049 so-far unevaluated Orchid species in a short time and increases the proportion of evaluated species by 53% across geographic regions. IUC-NN was robust to data availability, was less biased than the IUCN Red List and AA were robust to errors in the geographic input data (question 3).

## Supporting information

Supplementary Material S3 - Conservation assessments for 13,910 Orchid species

Supplementary Material S4 - The impact of feature choice on IUC-NN

Supplementary Material S5 - Per species comparison of IUC-NN and IUCN RL

## Supporting Information

Supplementary Material S1 - Supplementary Methods

Supplementary Material S2 - Supplementary Tables and Figures

Supplementary Material S3 - Conservation assessments for 13,910 Orchid species

Supplementary Material S4 - The impact of feature choice on IUC-NN

Supplementary Material S5 - Per species comparison of IUC-NN and IUCN RL

## Acknowledgements

We thank WCSP for provision of the species list and information on life form and geographic distribution, as well as all data contributors and publishers contributing to GBIF for their effort to collect, digitize, store and publish orchid records. We thank A. T. Clark for discussion and help on the permutation test. AZ and PV acknowledge funding by sDiv, the Synthesis Center for Biodiversity Sciences - a unit of the German Centre for Integrative Biodiversity Research (iDiv) Halle-Jena-Leipzig, funded by the German Research Foundation (FZT 118). TMK acknowledges funding from the Alexander von Humboldt Foundation in the framework of the Alexander von Humboldt Professorship. DS received funding from the Swiss National Science Foundation (PCEFP3_187012; FN-1749) and from the Swedish Research Council (VR: 2019-04739).

