## Supplementary Material S1 - Supplementary Methods for "Automated conservation assessment of the orchid family using deep learning"

#### Automated assessmnets using index-based methods

We generated AAs using three index-based methods: ConR v.1.2.2 (Dauby et al. 2017), speciesgeocodeR v.2.0-10 (Schmidt et al. 2017; Töpel et al. 2017) and rCAT v.0.1.5 (Moat 2017). ConR calculates three metrics following IUCN Criterion B: the extent of occurrence (EOO), the area of occupancy (AOO), and the number of locations. Together with the assumption of declining habitat quality/range size for all species, ConR uses these indices to classify species into IUCN threat categories as B1ab(i/ii/iii/iv/v), B2ab(i/ii/iii/iv/v), or B1ab(i/ii/iii/iv/v)+2ab(i/ii/iii/iv/v). We ran ConR using the `IUCN.eval` function, with a cell size of 2x2 sqkm for the calculation of AOO, as recommended by the IUCN (IUCN Standards and Petitions Subcommittee 2017) and a cell size of 10x10 km for estimating the number of locations as suggested by the developers of ConR (Dauby et al. 2017). We did not include protected areas in the analyses since there is a high uncertainty associated with the efficiency of different protected areas, i.e. "paper parks". The rCAT and speciesgeocodeR methods work similarly to ConR except that they only use EOO and AOO.

We then used existing IUCN Red List assessments as standard and compared the overall accuracy of these methods to assess species in the same category as IUCN. We did so on two levels: a detailed level, including four categories (*CR*, *EN*, *VU*, *NT or LC*) as well as on a binary *Possibly Threatened* (*CR*, *EN*, *VU*) *vs. Not Threatened* (*LC or NT*) level. We provide full error matrices for both cases. For the binary assessment, we also calculated the fraction of false positives (non-threatened species classified as *Possibly Threatened* by AA) and false negatives (threatened species classified as *Not Threatened* by AA).

#### Automated assessments using IUC-NN

In a second step, we developed a new prediction-based method—IUC-NN—based on a deep neural network (Goodfellow et al. 2016). IUC-NN uses species with existing IUCN assessments to train a neural network to predict the conservation status of species so far not evaluated. We based the training and prediction on a set of per species traits (features), which we derived from the geographic occurrences for each species, and we grouped the features into different subsets.

The *geographic features* included mean latitude of the occurrence records, mean longitude, latitudinal range, longitudinal range, the number of occurrence records available, the EOO, the AOO, and the number of locations (the last three obtained from ConR from step 1). In addition we obtained 19 bioclim variables (*climate features*; Karger et al. 2017), biome information (*biome features*; Olson et al. 2001) and the human footprint index (Wildlife Conservation Society - WCS & International Earth Science Information Network - CIESIN - Columbia University 2005; Di Marco & Santini 2015; Darrah et al. 2017) for all records and averaged them for each species. We restricted our analyses to traits we could derive from geographic occurrences, since our aim was to test how well digitally available occurrences can inform conservation assessment, and because additional data was only available for few species. To test the importance of different predictor types, we tested different combinations of groups of features, as well as all features combined. IUC-NN is implemented in Python (v. 3.7) using the `TensorFlow` (https://www.tensorflow.org) platform.

We tested two classification schemes: one with two classes (*Possibly Threatened* vs. *Not Threatened* comparable to the ConR results) and one with all five IUCN classes (*CR*, *EN*, *VU*, *NT*, and *LC*). Prior to the training, we randomly split the dataset into a training and validation set (90% of the entries) and a test set (10%). To carry out the training we further split the dataset in 80% of the entries for training and 20% for validation.

As the size of the dataset was comparatively small, we performed cross-validation by shifting five times the validation set to quantify the average validation cross-entropy loss (a measure of the distance between predicted and true values) and the average validation accuracy.

Across all analyses, we used a "Glorot normal" kernel initializer with the "ReLU" activation function for the hidden layers and a "SoftMax" function to obtain probabilities associated with each class (Goodfellow et al. 2016). We set the number of nodes equal to the number of features with the default inclusion of the bias vector (https://keras.io). After running the first training for 5,000 epochs, we computed the number of epochs that yielded the lowest cross-entropy loss. We then averaged the number of epochs across the five replicates and re-trained the network this time using the entire data set (excluding the test set). We then used the re-trained network to compute the test accuracy.

We repeated the protocol described above for different numbers of hidden layers ranging from one to five to determine the architecture the resulted in the lowest cross-entropy loss. Additionally, we repeated the analyses using configuration of nodes and layers but applying Dropout with probability of 0.3 in the first layer (Srivastava et al. 2014). We chose as the best neural network the one yielding the lowest cross-validation cross-entropy loss and used it to estimate the test accuracy and assess the status of all orchid species lacking an IUCN assessment. We also used it to compute confusion matrices from the entire set of species assessed by IUCN for comparison with the other methods. The IUC-NN method is implemented in the Python program `ruNNer` (https://github.com/dsilvestro/ruNNer) and the trained neural networks used for the empirical analyses shown here are available as Supplementary Material S1.

### Sensitivity analyses

#### The reliability of automated assessments

##### Data availability

To address **question 2**, we first tested how the number of records available affects the accuracy of AA methods. To do so, we fitted four separate binomial regression models to estimate the relationship between the AA of a species agreeing with the IUCN RL status and the number of occurrence records available for this species, using the `stats::glm` function in R, with a logit link. We created one model for speciesgeocoder, rCAT, ConR, and IUC-NN, respectively (n = 861 species each). For IUC-NN this included the predictions for training, test and validation data.

##### Data quality

Since it is unclear if geographic errors in the occurrence information bias automated assessments (Zizka et al. 2020b), we repeated the benchmarking of the index-based AA based on different datasets of species occurrences, representing three levels of data curation (1) the data downloaded from GBIF scrubbed taxonomically ("raw" hereafter). (2) The raw dataset excluding geographic errors common to public databases (Zizka et al. 2019), excluding i) records with incomplete names, and restricting the data set to accepted names from the WCSP, ii) records without geographic coordinates; iii) records older than 1900, since they are often imprecise due to *post-hoc* geo-referencing from vague locality descriptions; iv) records based on fossils, tissue samples or living collections; v) records with a reported uncertainty higher than 100 km; vi) records in the sea, assigned to country or province centroids, assigned to the location of biodiversity institutions, zero coordinates, and equal latitude and longitude as flagged by the 'CoordinateCleaner' v. 2.0-9 R package (Zizka et al. 2019). Since coordinates assigned to country centroids are a well-known problem (Maldonado et al. 2015) we additionally checked individual localities with many records in the data set and excluded all records where the locality was the centroid of a country, and vii) multiple records of species from individual localities, in which case we only used one ("intermediate", hereafter). And (3) the intermediate dataset with additional removal of records outside the registered botanical country if data from the WCSP (WCSP 2019) was available ("filtered", hereafter). For all datasets, we rounded coordinates to four decimals. We then compared the overall accuracy of ConR for all datasets. We only tested the effect of data quality on AA for ConR, since we expect the two other index-based methods to respond similarly to the issue and because we expect IUC-NN to be more robust against erroneous coordinates, since the features used for prediction are mean values across all records of a species.

##### Sampling bias

Since the collection of geographic records and the evaluation process for the IUCN RL may be biased by geographic region or life form (Meyer et al. 2016; Daru et al. 2018; Zizka et al. 2020a), we tested the biasing effect of life form and geographic distribution on both AA and IUCN Red Listing. To do so, we classified species as 'epiphyte', 'lithophyte', 'terrestrial' or 'mixed' (species found growing as both epiphytes/lithophytes and/or terrestrial) based on the WCSP life form data. Then we calculated the fraction of each life form of (1) all orchid species with known life form (n = 24,204 species), (2) species with known life form evaluated by AA (n = 12,118 species), and (3) species with known life form evaluated by the IUCN RL (n = 907 species). We used a similar classification for continents, assigning each species to continental regions based on the SCSP distributions information (TDWG level-1), and calculated the fraction of species in each region for 1) all species with known distribution (n = 26,588 species), 2) species evaluated by AA (n = 13,910 species), and 3) species evaluated by the IUCN Red list (n = 1,195 species) and compared the fractions as for life form. We then tested whether observed differences among assessments were greater than expected by random chance based on a null model in which we randomly permuted bias measurements among model types.

##### Alternative reference standards

Because the IUCN RL is biased geographically and taxonomically (www.iucn.org), we repeated the accuracy tests of the index-based AA with four additional reference datasets: (1) assessments available from the sampled red list index ("SRLI", hereafter) which is a random sub sample of all available IUCN assessments to reduce taxonomic bias (Royal Botanic Gardens 2010); (2) a subset of the IUCN assessments including those species assessed after 2008, to ensure up-to-date assessments and a standardized methodology; (3) a subset of the IUCN assessments including species with at least 15 occurrence records available for AA, since this number has been suggested as minimum for reliable AA (Rivers et al. 2011); and (4) an additional dataset of species conservation assessments from the ThreatSearch database, which gathers conservation assessments from the literature. In the case of the ThreatSearch database, we only retained global assessments and ignored species assessed as *Data Deficient*. We only tested these alternative references with ConR, since we expect the other index-based methods to have similar results, and the alternative datasets included too few species to train IUC-NN models.

#### The status and global hot spots of Orchid extinction risk

To address **question 3** we used IUC-NN for an AA of all orchid species for which we retained occurrence records after full geographic filtering. We chose IUC-NN, since this method was most accurate, and chose as the best neural network the one yielding the lowest cross-validation cross-entropy loss and used it to assess the status of all orchid species lacking an IUCN assessment.

We quantified the species coverage of IUC-NN in comparison to IUCN-RL, by identifying the proportion of global orchid species evaluated with both methods. We used the AA obtained from IUC-NN together with the geographic distribution from the WCSP to visualize the number and fraction of *Possibly Threatened* species per botanical country (TDWG level 3). Furthermore, we summarized the number of orchid species and the fraction of threatened species in major biomes of the world. We classified species into biomes based on a global biome scheme (Olson et al. 2001) and on their occurrence records, using the 'speciesgeocodeR' v. 2.0-10 package (Töpel et al. 2017), if at least five percent of a species' records occurred in this biome (see Antonelli et al. 2018 for a justification of the threshold choice). To indicate the uncertainty of our global assessment, related to the number of not evaluated species, we provide an estimate of the number of missing species per bioregion.
