## Supplementary Material S2 - Supplementary Tables and Figures for "Automated conservation assessment of the orchid family using deep learning"

Table 1: The impact of individual geographic filters on the number of occurrence records and taxa available.

| Filter | Occurences remaining after filtering | Taxa remaining after filtering |
|---|---|---|
| Records without coordinates | 3,050,875 | 17,971 |
| Records older than 1900 | 2,972,297 | 17,790 |
| Based on other data than specimen, observation and literature | 2,938,432 | 17,583 |
| Records with less than one or more than 99 individuals | 2,907,504 | 17,571 |
| Precision below 100km | 2,906,487 | 17,546 |
| Duplicated information | 1,215,545 | 17,546 |
| Automatic filters: capitals, centroids of countries and provinces, equal lon/lat, GBIF headquarters, the sea, biodiversity institutions or plain zero longitude or latitude | 1,205,984 | 17,238 |
| Additional country centroids | 1,201,316 | 16,935 |
| Duplicates after rounding coordinates to four decimals | 1,188,658 | 16,935 |
| Records disagreeing with distribution information on TDWG region | 999,476 | 14,148 |

Table 2: The confusion matrices for index-based automated conservation assessment methods compared to conservation assessments provided on the IUCN RL in four detailed categories.

| | | IUCN Assessment | | | |
|---|---|---|---|---|---|
| AA method | Spatial cleaning | CR | EN | VU | LC or NT |
| **ConR** | | | | | |
| CR | full | 31.4 | 26.1 | 10.5 | 32.0 |
| EN | full | 11.3 | 36.7 | 12.3 | 39.7 |
| VU | full | 2.7 | 35.1 | 18.2 | 43.9 |
| LC or NT | full | 0.8 | 8.3 | 15.8 | 75.1 |
| CR | medium | 29.1 | 27.3 | 12.7 | 30.9 |
| EN | medium | 13.1 | 38.5 | 11.2 | 37.2 |
| VU | medium | 3.1 | 34.4 | 22.1 | 40.5 |
| LC or NT | medium | 1.1 | 8.7 | 15.2 | 75.1 |
| CR | raw | 36.6 | 24.2 | 11.1 | 28.1 |
| EN | raw | 15.0 | 39.8 | 9.8 | 35.5 |
| VU | raw | 2.8 | 32.8 | 23.2 | 41.2 |
| LC or NT | raw | 0.7 | 10.3 | 15.2 | 73.8 |
| **rCat** | | | | | |
| CR | full | 27.6 | 28.3 | 9.4 | 34.6 |
| EN | full | 8.1 | 56.6 | 12.1 | 23.2 |
| VU | full | 2.2 | 39.8 | 21.5 | 36.6 |
| LC or NT | full | 1.9 | 14.0 | 15.7 | 68.3 |
| **SPGC** | | | | | |
| CR | full | 42.9 | 42.9 | 0.0 | 14.3 |
| EN | full | 21.7 | 36.1 | 10.4 | 31.8 |
| VU | full | 2.1 | 40.4 | 21.3 | 36.2 |
| LC or NT | full | 1.9 | 13.8 | 15.8 | 68.5 |

Table 3: The confusion matrix for the prediction based IUC-NN model predicting detailed IUCN RL classes.

| | IUCN assessment | | | | |
|---|---|---|---|---|---|
| IUC-NN | CR | EN | VU | NT | LC |
| **Best Model** | | | | | |
| CR | 46.2 | 30.8 | 0.0 | 0 | 23.1 |
| EN | 4.2 | 83.3 | 4.2 | 0 | 8.3 |
| VU | 0.0 | 57.1 | 0.0 | 0 | 42.9 |
| NT | 0.0 | 0.0 | 0.0 | 25 | 75.0 |
| LC | 0.0 | 11.8 | 0.0 | 0 | 88.2 |
| **All features** | | | | | |
| CR | 38.5 | 46.2 | 0.0 | 0 | 15.4 |
| EN | 0.0 | 87.5 | 0.0 | 0 | 12.5 |
| VU | 7.1 | 35.7 | 14.3 | 0 | 42.9 |
| NT | 0.0 | 50.0 | 0.0 | 0 | 50.0 |
| LC | 2.9 | 11.8 | 0.0 | 0 | 85.3 |

Table 4: The confusion matrix for ConR using different data than the global IUCN RL as gold standard.

| | IUCN Assessment | |
|---|---|---|
| AA method | Not Threatened | Possibly Threatened |
| **IUCN, last 10 years** | | |
| Not Threatened | 77.6 | 22.4 |
| Possibly Threatened | 41.6 | 58.4 |
| **IUCN > 15 records** | | |
| Not Threatened | 75.3 | 24.7 |
| Possibly Threatened | 44.8 | 55.2 |
| **Sampled Red List Index** | | |
| Not Threatened | 96.1 | 3.9 |
| Possibly Threatened | 67.9 | 32.1 |
| **ThreatSearch** | | |
| Not Threatened | 62.5 | 37.5 |
| Possibly Threatened | 0.0 | 100.0 |

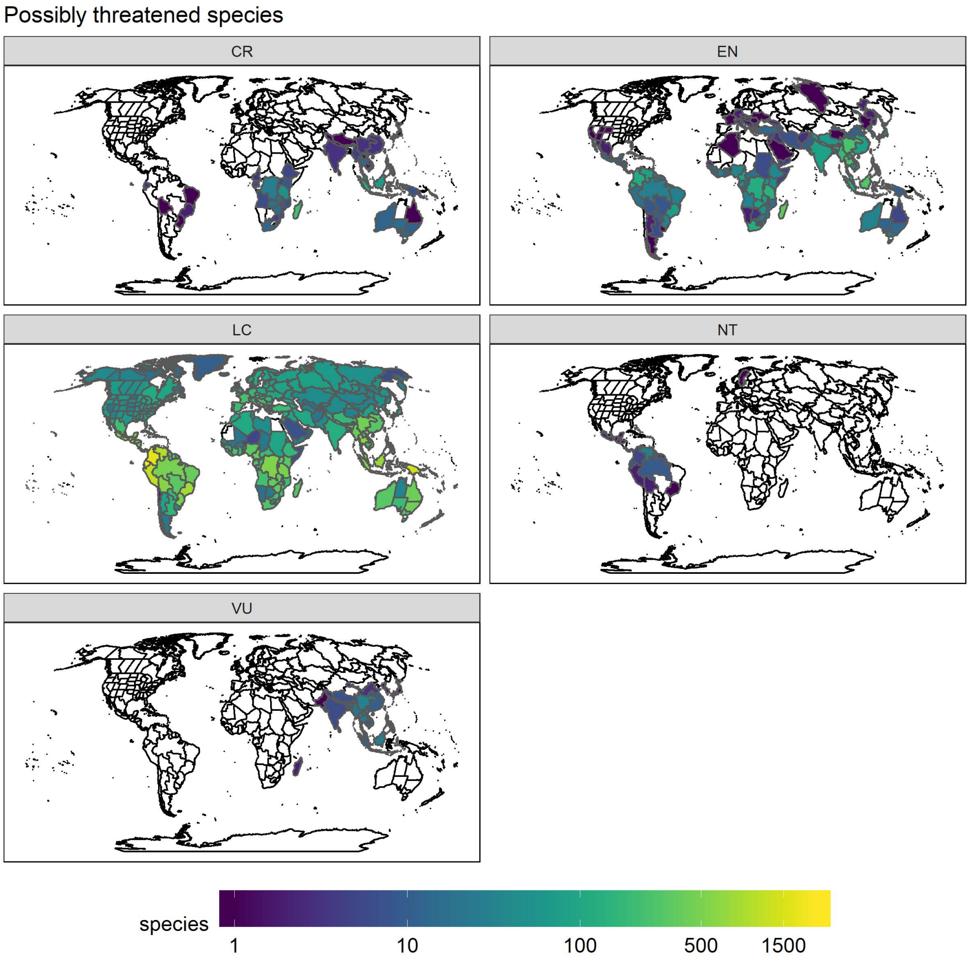


Figure 1: The number of orchid species classified as *Critically Endangered (CR)*, *Endangered (EN)*, *Vulnerable (VU)*, *Near Threatened (NT)*, and *Least Concern (LC)* by the automated conservation assessment using IUC-NN per botanical country.

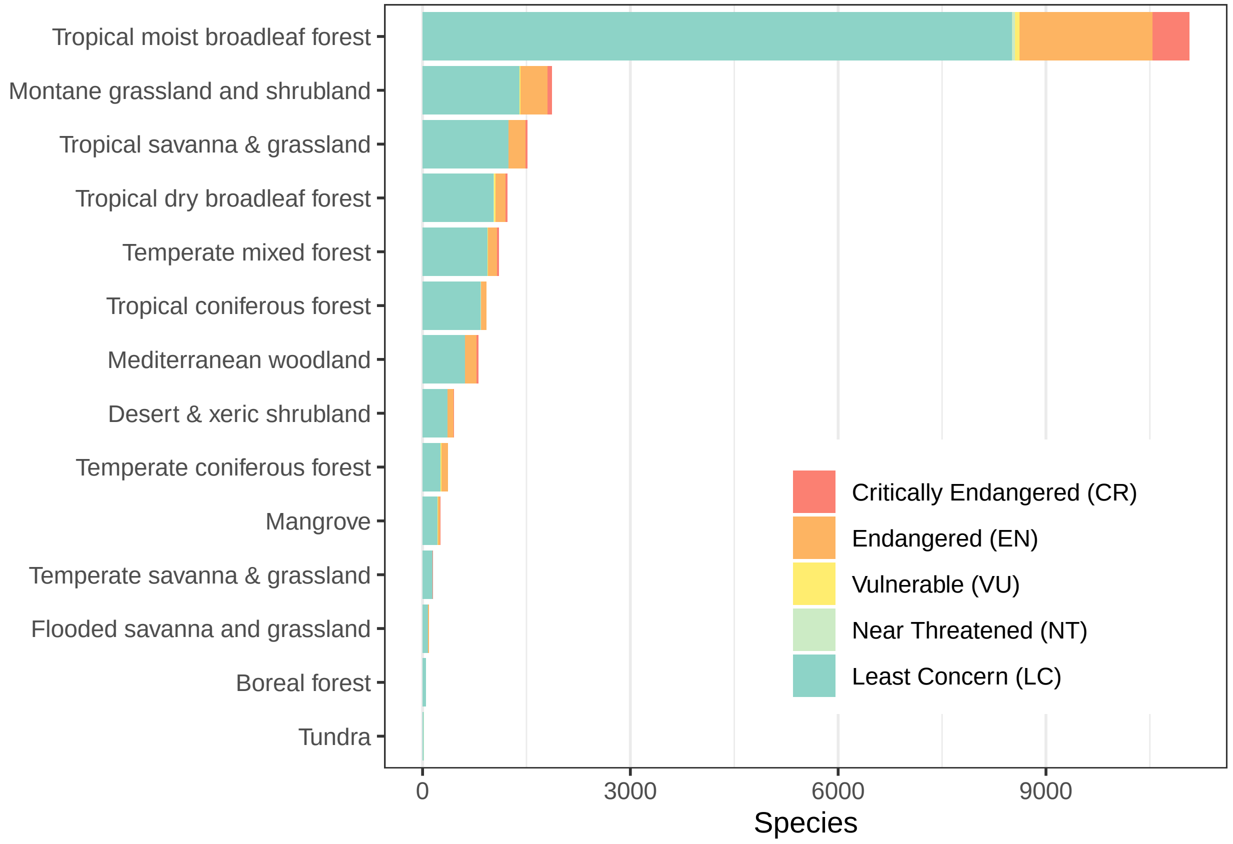


Figure 2: Detailed conservation status of orchids per biome, based on the automated assessment. Biomes following Olson et al. (2001), the biome names have been shortened for better readability.

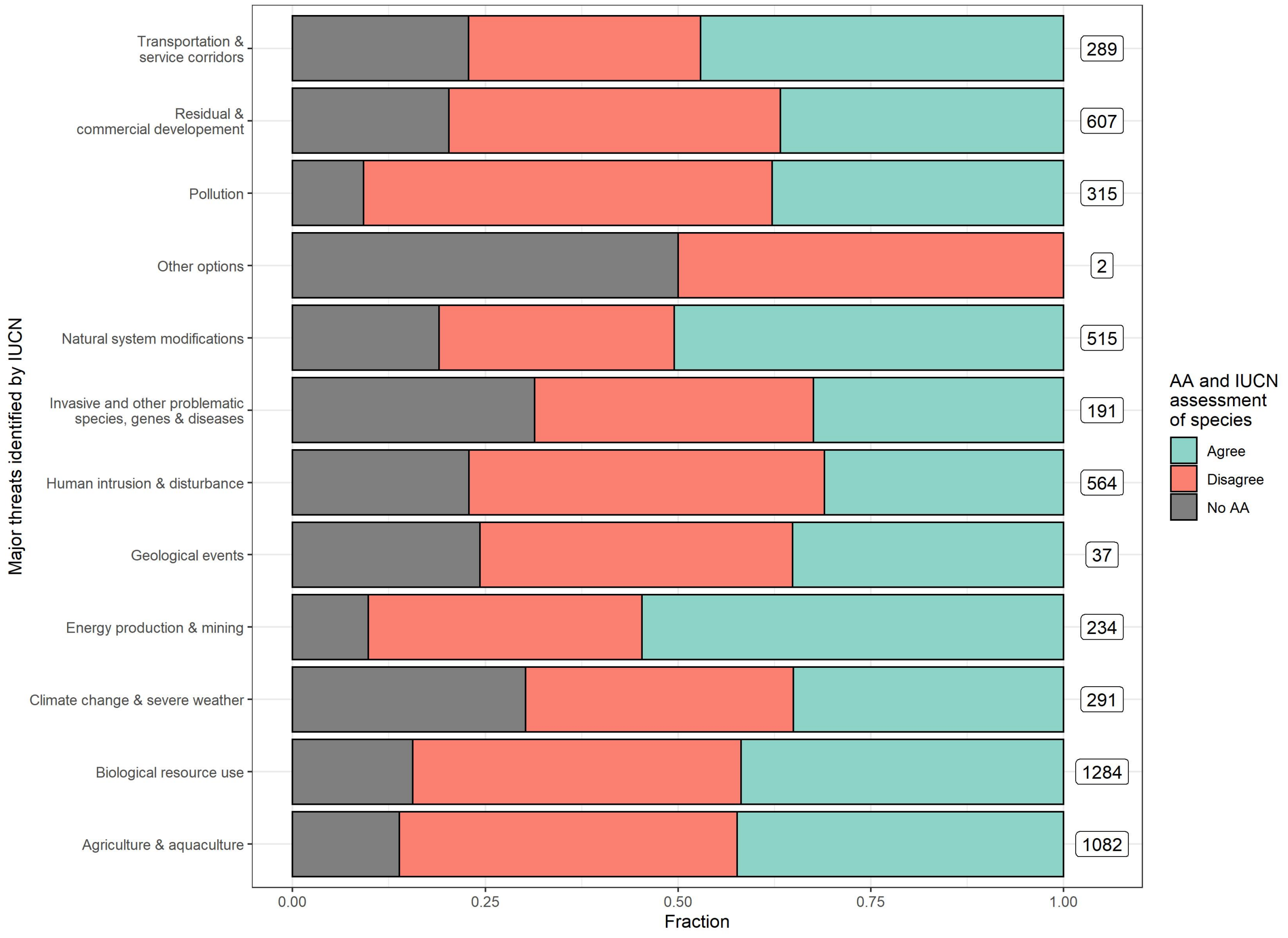


Figure 3: The relationship between prediction accuracy of automated assessments and the threat categories identified by IUCN RL for a species.

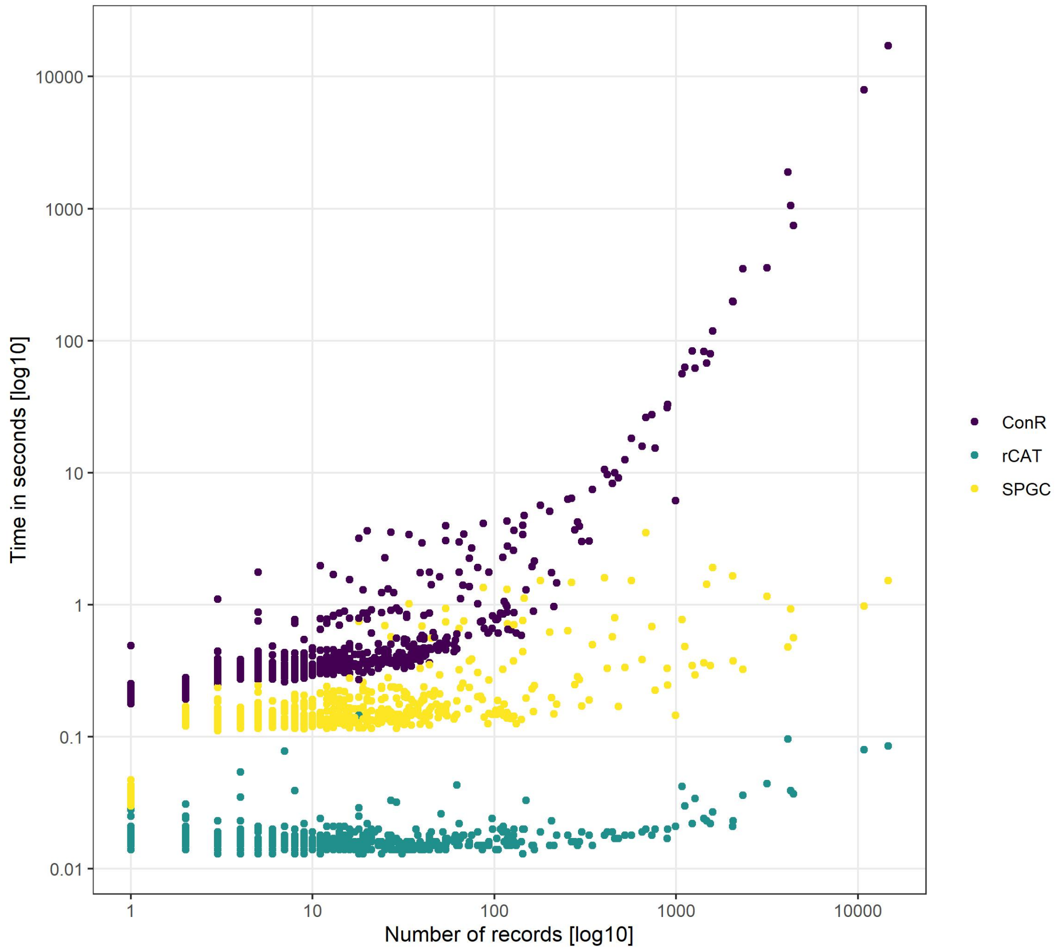


Figure 4: The computation time necessary for index-based AA with different methods. Computation time increases with the number of records, most importantly so for the ConR method. Note the log transformation of both axes. Most species were evaluated in less than one second.

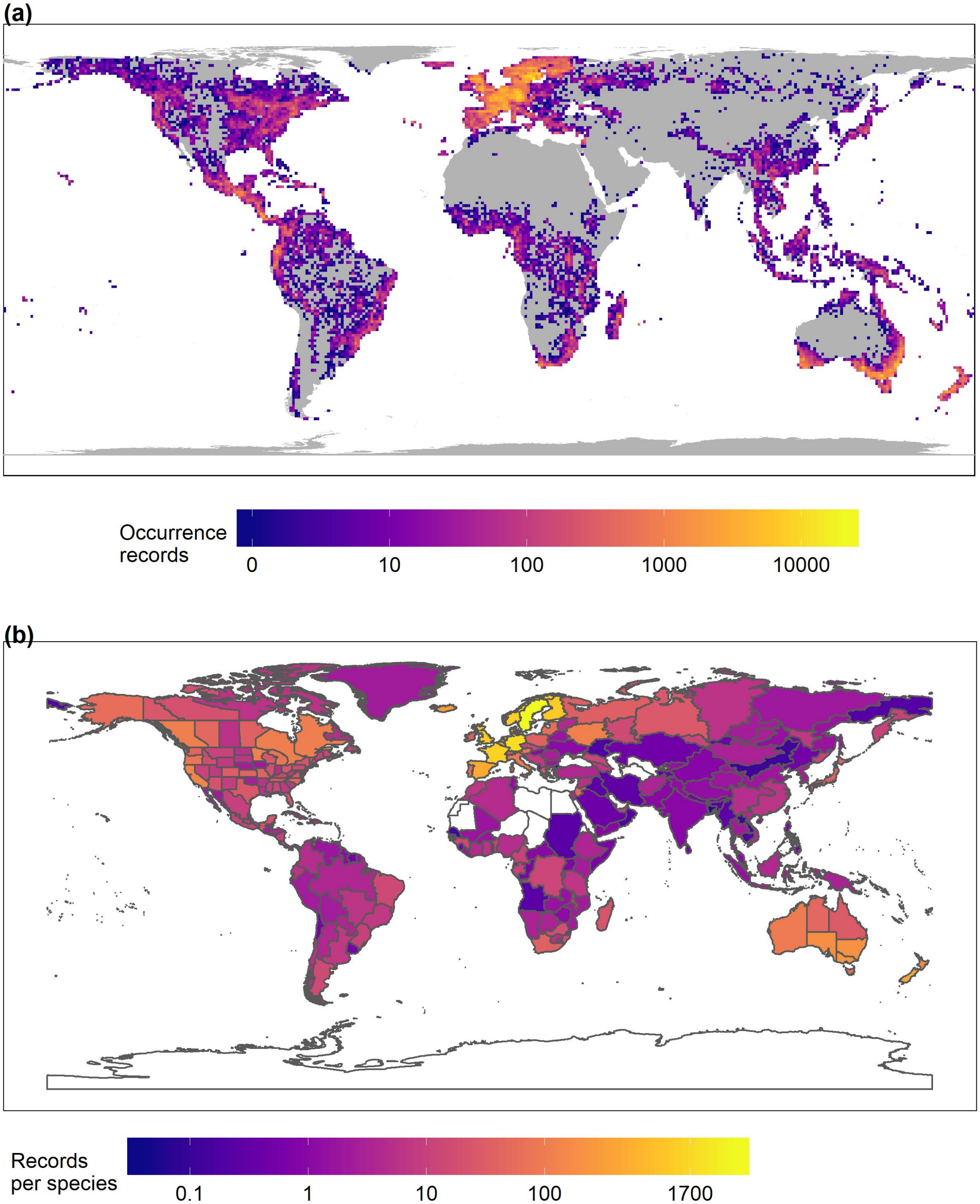


Figure 5: The geographic sampling used in this study. A) The number of orchid records remaining after filtering, in a 100x100 km grid. B) The number of records per orchid species available after filtering in a given TDWG region. Sampling is biased towards certain regions, especially Central Europe.

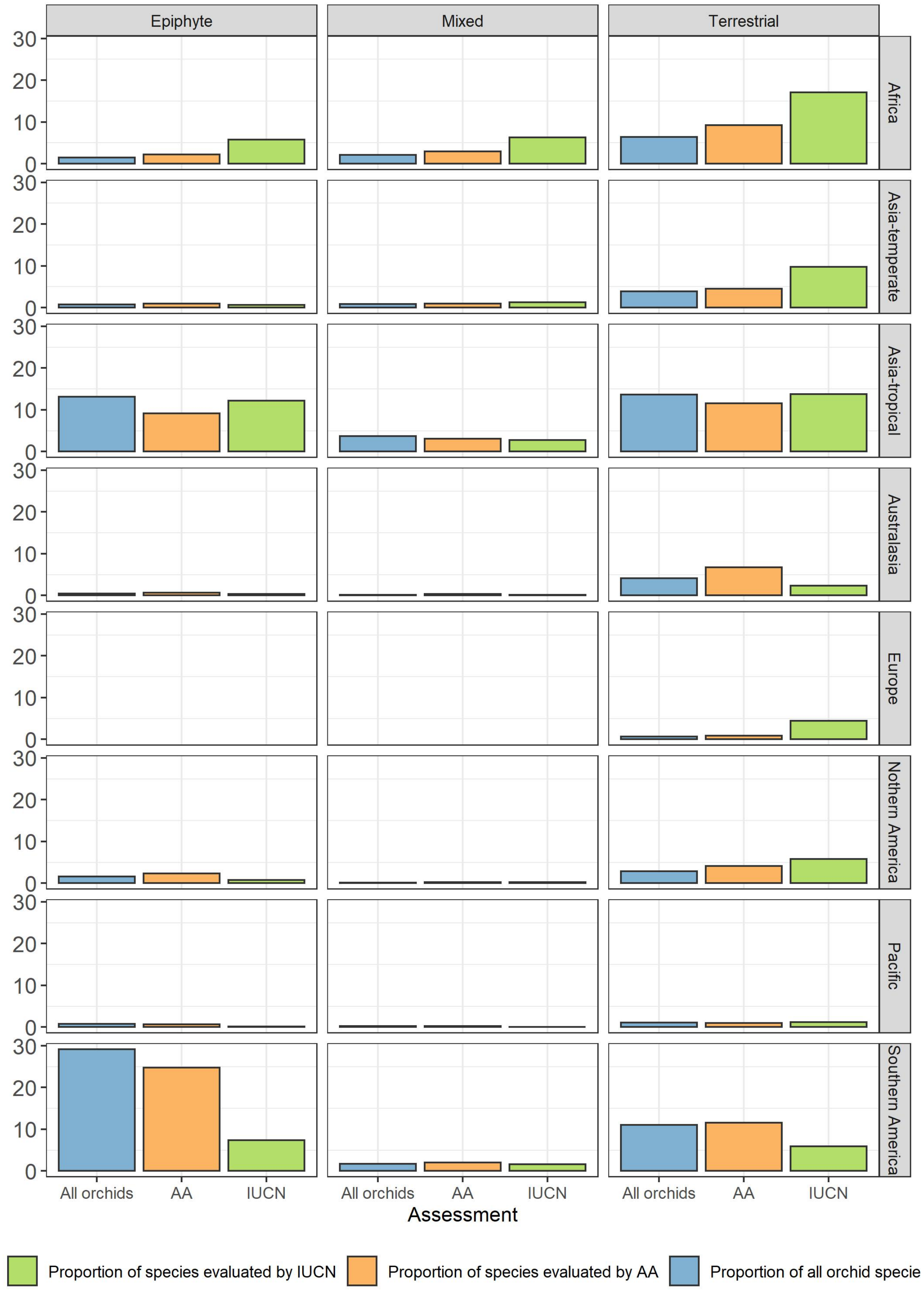


Figure 6: The proportion of species represented in each life form class and each continental region (TDWG level 1), for three datasets: all orchid species with life form and geographic data available (blue), all species evaluated using automated assessment (orange), and all species with an IUCN RL assessment available.
